# Species boundaries, host associations, and possible host shifts in the turtle barnacle genus *Platylepas*

**DOI:** 10.1101/2025.09.03.674014

**Authors:** Ryota Hayash, Hiroyuki Ozawa, Takashi Seiko, Takahide Sasai, Takushi Kishida

## Abstract

The barnacle genus *Platylepas* contains obligate epibionts of marine vertebrates, but the taxonomic status and evolutionary history of the sea-snake specialist, *P. ophiophila*, have remained unclear. Here, we reassess species boundaries and host associations in *Platylepas* using shell-morphological comparisons and sequence data from mitochondrial COI and nuclear H3 genes from specimens collected from sea turtles, sea snakes, and a dugong. Our results show that *P. ophiophila* is a valid species, phylogenetically and morphologically distinct from the widespread turtle barnacles *P. decorata* and *P. hexastylos*. Furthermore, we report the first record of *P. ophiophila* from a sirenian host (*Dugong dugon*), a discovery that necessitates the re-examination of historical records of barnacles on sirenians worldwide. Our phylogeny places *P. decorata*, a turtle epibiont, as the earliest diverging lineage within the genus. This is consistent with the hypothesis that the ancestor of *Platylepas* was associated with sea turtles, a conclusion supported by the ichnofossil record. The subsequent divergence of the *P. ophiophila* clade may represent an important host-use transition: an ancient host shift from sea turtles to the taxonomically disparate but ecologically linked inhabitants—sea snakes and sirenians—of seagrass ecosystems. This study clarifies species limits within *Platylepas* and provides a framework for discussing host associations and potential host shifts in light of existing molecular and fossil evidence.

## INTRODUCTION

Barnacles of the superfamily Coronuloidea are obligate epibionts on marine vertebrates, including sea turtles, cetaceans, and sea snakes (Newman and Ross 1976, Hayashi 2013a, 2025). Among these, species of the genus *Platylepas* exhibit diverse host associations and complex morphological variation, often complicating their taxonomic resolution. Taxonomically and morphologically, *Platylepas* is highly specialized for an embedded lifestyle, which sharply contrasts with the attachment strategy of its close relative, *Chelonibia*. While *Chelonibia* species primarily attach superficially to hard substrates (e.g. the carapace of sea turtles) by cementation, *Platylepas* secures itself by becoming partially embedded in the host epidermis or scutes and mechanically interlocking with the surrounding tissue. Morphologically, *Platylepas* possesses a fragile, six-plated shell characterized by a membranous basis and prominent internal midribs on each compartment (Hayashi, 2012). These midribs act as structural anchors that push deep into the relatively softer epidermis or scutes of the host, allowing the host’s growing tissue to envelop the barnacle shell. This interlocking mechanism provides robust physical retention on flexible or frequently shedding surfaces, such as the skin and scutes of sea turtles.

Historically, the genus *Platylepas* was established by Gray (1825) based on these unique morphological adaptations. Since its original description from sea turtles, subsequent marine biological surveys progressively revealed a much broader host range, expanding to include sea snakes and sirenians. However, this diverse host utilization has been a major source of taxonomic controversy. Because the shell morphology of *Platylepas* is known to be influenced by the physical properties of the host substrate, it has historically been difficult to determine whether morphological variations represent distinct host-specific species or merely phenotypic plasticity (ecotypes) within a single widespread taxon. A striking precedent for such taxonomic overestimation exists in the closely related genus *Chelonibia*. In this group, molecular analyses have revealed that nominal species previously distinguished by host association and shell morphology—such as the turtle-associated *C. testudinaria*, the crustacean-associated *C. patula*, and the sirenian-associated *C. manati*—actually represent a single, highly plastic species (Cheang et al., 2013; Zardus et al., 2014).

To understand the evolutionary context of this highly specialized host-epibiont relationship, both molecular and paleontological data must be considered. Molecular phylogenetic analyses indicate that the superfamily Coronuloidea, including the turtle barnacle lineages, originated and diversified during the Miocene (Hayashi et al., 2013). Historically, a temporal discrepancy existed between these molecular estimates and the paleontological data, as the body fossil record of *Platylepas* is notably sparse and predominantly restricted to the Pleistocene, such as *P. hexastylos* from the Atsumi peninsula, Japan (Karasawa and Kobayashi, 2022), the extinct †*P. wilsoni* from Florida, USA (Ross, 1963), †*P. mediterranea* from the Mediterranean basin (Collareta et al., 2019), and an unidentified fossil *Platylepas* sp. from the Hirano formation, Japan (Mimoto, 1991). Recently, however, this gap has been reconciled by ichnological evidence. Prompted by the earlier suggestion that trace fossils on host remains could provide crucial clues to coronuloid evolutionary history (Hayashi et al., 2013), the discovery of diagnostic attachment scars (e.g., *Karethraichnus* borings) on Miocene turtle shells now provides robust fossil support for the ancient and deep-rooted association between platylepadid barnacles and their chelonian hosts (Collareta et al., 2022). Furthermore, recent fossil discoveries have been suggested to extend the temporal range of this highly specialized ’turtle-riding’ lifestyle even further back to the early Oligocene, implying that the symbiotic association between coronuloid barnacles and sea turtles may have persisted for over 30 million years (Collareta et al., 2023).

While *Platylepas hexastylos* and *P. decorata* are frequently reported from sea turtles, *P. ophiophila* has been described primarily from sea snakes and is considered a putative sea-snake specialist, and its taxonomic status remains uncertain. Drawing parallels to the situation in *Chelonibia* (Cheang et al., 2013; Zardus et al., 2014), some researchers have suggested that it may represent a morphological variant or ecotype of *P. hexastylos*, based on overlapping traits and limited molecular data (Pfaller et al. 2012). Clarifying the species boundaries within *Platylepas* is essential not only for barnacle taxonomy but also for understanding host specificity and epibiotic relationships in marine ecosystems.

In this study, we examine *Platylepas* specimens collected from sea turtles (*Caretta caretta*, *Chelonia mydas*, *Eretmochelys imbricata*), sea snakes (*Hydrophis melanocephalus*) and a dugong (*Dugong dugon*) in Okinawa, Japan. Using shell-morphological comparisons and COI/H3 sequence data, we reassess the distinctiveness of *P. ophiophila* from *P. hexastylos*, evaluate the identity of dugong-associated *Platylepas* specimens, and discuss the implications of these findings for host associations, host-use evolution, and the use of epibiotic barnacles as indicators of host movement history.

Through detailed morphological comparisons and molecular phylogenetic analyses of mitochondrial COI and nuclear H3 genes, we aim to reassess the validity of *P. ophiophila* and evaluate its distinction from *P. hexastylos*. Furthermore, this study presents the first record of *P. ophiophila* from a sirenian host, a discovery that broadens our understanding of its host range and the phylogenetic relationships within the genus *Platylepas*.

## MATERIALS AND METHODS

### Field sampling

Epibiotic surveys of marine vertebrates were conducted on the main island of Okinawa, subtropical area of Japan (Fig. 1, Table 1). Slender-necked sea snakes (*Hydrophis melanocephalus*) were collected in Awase and Motobu, and a dugong (*Dugong dugon*) was stranded in Nakijin, Okinawa on 18 March 2019, and an autopsy was conducted on 17 July 2019 (Okinawa Prefecture Government 2021, Ozawa *et al*. 2024). For comparative purposes, three species of sea turtles (*Caretta caretta*, *Chelonia mydas*, *Eretomochelys imbricata*) were also examined for epibionts.

**Figure 1.**
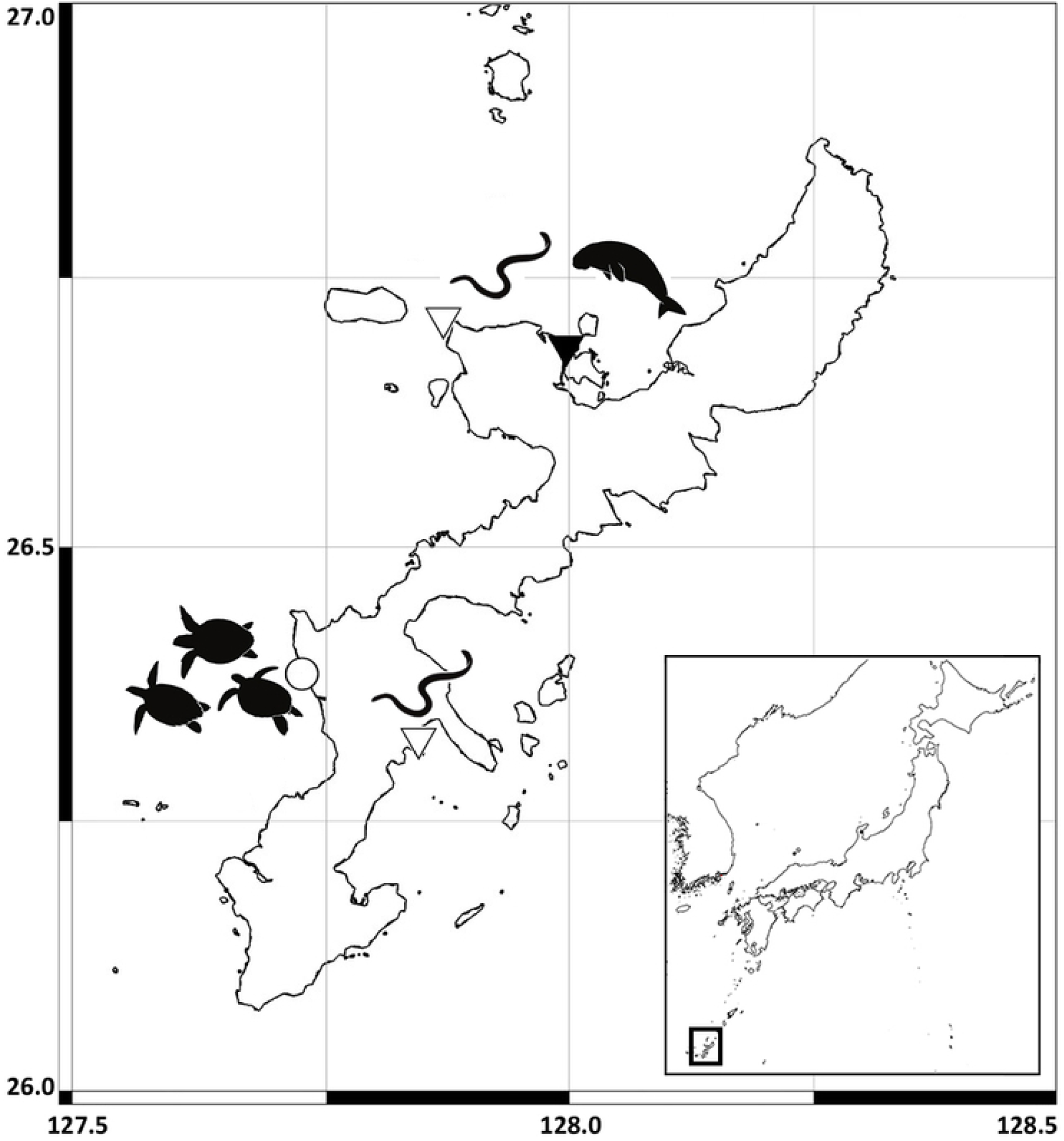
Geographical locations of sampling sites for a dugong (black triangle), sea snakes (white triangles), and sea turtles (circle).

**Table 1.** Materials examined and GenBank accession number of the sequences. *Histone 3 sequences of *Platylepas decorata* and *Chelonibia caretta* were used from Hayashi et al. (2013), AB723941 and AB723933.

Barnacles were removed from hosts and preserved in 99% ethanol.

### Morphological examination

Specimens were examined under a stereomicroscope. Shell outline, parietal ornamentation, midrib development, basis, and opercular plates were compared with previous descriptions and illustrations of *Platylepas* species. The specimens were identified in the laboratory and subsequently deposited in National Museum of Natural Science, Tokyo (NMNS-Cr-33017-33026).

### DNA extraction, sequencing, and Phylogenetic analysis

DNA was extracted from muscle tissue using the commercial kit (Tissue Genomic DNA Extraction Micro Elute Kit, Favorgen). Partial sequences of mitochondrial and nuclear genes were amplified the PCR primer set for the COI gene (LCO-1490 / B2R-COI) and the Histone 3 gene (AF / AR) (Folmer *et al*. 1994, Ayre *et al*. 2009, Colgan *et al*. 1998). PCR was performed in a 10 μl of a reaction volume containing 5.0 μl KOD One DNA Polymerase (Toyobo), 3.2 μl deuterium-depleted water, 0.5 μl of each primer (5 μM) and 1.0 μl of DNA template.

The thermal regime entailed initialization using 35 cycles for 10 seconds for denaturation at 98°C, 5 seconds for annealing at 48°C for COI and 50 °C for H3, and 1 second for extension at 68 °C, and final extension at 15°C. The PCR products were observed on 1.0% agarose gel, and the most intense products were used for Sanger sequencing. PCR products were purified once more using the Big Dye Xterminator kit (Thermofisher) and then sequenced on an ABI 3130 capillary sequencer. All sequences were aligned in MEGA X (Kumar *et al*. 2018) with MUSCLE (Edgar 2004) and trimmed to 5-600 bp for final alignment. All sequences were deposited in GenBank (Table 1).

## RESULTS

### Nomenclatural considerations: Platylepas ophiophilus or Platylepas ophiophila?

The sea snake barnacle was originally described as *Platylepas ophiophilus* by Lanchester (1902). The specific epithet is derived from the Greek *ophio-* (snake) and the Latinized suffix *-philus* (loving), forming a Latinized adjective meaning ‘snake-loving’. According to the International Code of Zoological Nomenclature (ICZN 1999), specifically Articles 31.2 and 34.2, a species-group name that is an adjective in the nominative singular must agree in gender with the generic name with which it is combined.

The genus name *Platylepas* is feminine. Consequently, the masculine adjectival epithet *ophiophilus* must be emended to the feminine form *ophiophila*. While the original spelling *ophiophilus* has persisted in various ecological and taxonomic literature (e.g., Zann 1975, Pfaller et al. 2012), the gender-conforming spelling *P. ophiophila* has been correctly applied by several authors in the past (e.g., Utinomi 1970, Ren 1980, Buzás et al. 2018, Collareta et al. 2019, Hayashi 2025). This mandatory emendation is strictly a spelling adjustment to satisfy grammatical gender agreement, not the proposal of an alternative species name; therefore, it fully aligns with the Code without threatening nomenclatural stability (Jiménez-Mejías et al. 2024). In accordance with these rules, we adopt the grammatically correct spelling *Platylepas ophiophila* throughout this study.

## SYSTEMATICS

Suborder **Balanomorpha** Pilsbry, 1916 Superfamily **Coronuloidea** Newman & Ross, 1976 Family **Platylepadidae** Newman & Ross, 1976 Genus ***Platylepas*** Gray, 1825 ***Platylepas decorata*** Darwin 1854 Fig. 2A-B.

**Figure 2.**
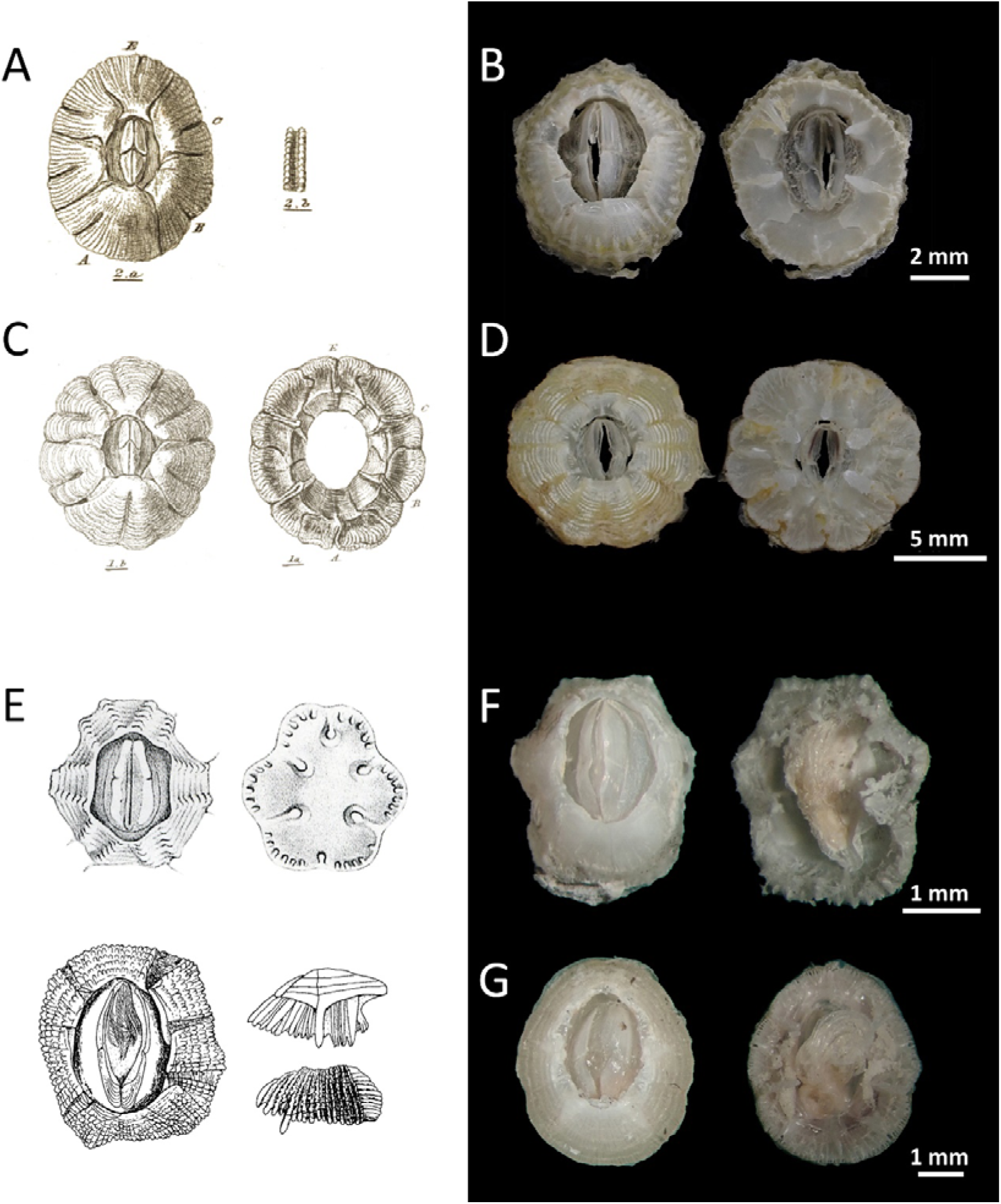
(A) *Platylepas decorata* reproduced from Darwin (1854); (B) *Platylepas decorata*, collected from a sea turtle, upper and basal views (NMNS-Cr-33017); (C) *Platylepas hexastylos* reproduced from Darwin (1854); (D) *Platylepas hexastylos*, collected from a sea turtle, upper and basal views (NMNS-Cr-33018); (E) *Platylepas ophiophila* reproduced from Lanchester (1902) and Utinomi (1970); (F) *Platylepas ophiophila*, collected from a black-headed sea snake, upper and basal views (NMNS-Cr-33023); (G) *Platylepas ophiophila*, collected from a dugong, upper and basal views (NMNS-Cr-33024).

*Platylepas decorata* Darwin, 1854: 429, pl. 17 figs. 2a-b. – Hayashi, 2012: 114, fig. 8; pl. 3b.

*Platylepas hexastylos ichtyophila* Pilsbry, 1916: 287, pl. 67 fig. 2. - Henry, 1954: 444.

*Platylepas multidecorata* Daniel, 1962: 641, figs. 1, 2.

### Descriptions

Shell subcircular, ring-like, shallow, steep sided, and solid; compartments with fine longitudinal ridges, lower edge dentate; each compartment with a midrib; radii narrow with simple septate margins; basal membrane equaling in convexity the shell; opercular valves oblong and narrow, scuta longer than terga.

### Previous records on Japanese coast

Ginoza, Okinawa (on *Chelonia mydas* and *Eretmochelys imbricata*); Ishigakijima Is., Okinawa (on *C. mydas* and *E. imbricata*); Ogimi, Okinawa (on *C. mydas*); Yomitan, Okinawa (on *C. mydas*); Yakushima Is., Kagoshima (on *C. mydas*); Hahajima Is., Ogasawara, Tokyo (on *C. mydas* and *E. imbricata*) (all records from Hayashi 2012).

### Habitat

Found on plastron, limbs and soft skin of sea turtles.

***Platylepas hexastylos*** (Fabricius, 1798) Fig. 2C-D

*Lepas hexastylos* Fabricius, 1798: 35, pl. 10 figs. 1, 2.

*Coronula bisexlobata* Blainville, 1824: 379, pl. 117 fig. 1.

#### *Platylepas pulchra* Gray 1825:105

*Platylepas bissexlobata* Darwin, 1854: 428, pl. 17 figs. 1a-d. - Gruvel, 1903:151, pl. 3 fig. 13.

*Platylepas hexastylos* Pilsbry, 1916: 285, pl. 67 figs. 1-1c, 3. – Hayashi, 2012: 114, fig. 7; pl. 3a.

*Platylepas hexast*y*los ichtyophila* Pilsbry, 1916: 287, pl. 67 fig. 2. - Young, 1991: 197, figs. 2g, h.

### Descriptions

Shell low, conical, margin multilobed, and orifice small; each compartments fimbriated at the edge, having a midrib; radii narrow; growth lines the predominant sculpturing, septae visible through the wall, becoming more numerous peripherally; line of ‘midrib fold’ obvious, dividing each compartment into two main lobes, accessory ‘folds’, and inter compartmental sutures, appear to enclose host tissue; midribs extend only slightly below level of periphery, forming a slightly convex base; septae visible on inner surface of outer wall towards periphery; opercular valves subequal, extending the full length of orifice.

#### Previous records on Japanese coast

Ginoza, Okinawa (on *Chelonia mydas* and *Caretta caretta*); Ohgimi, Okinawa (on *C. mydas*); Yomitan, Okinawa (on *C. mydas*, *Eretmochelys imbricata*, and *C. caretta* [Hayashi 2009]); Uruma, Okinawa (on *E. imbricata*); Ishigakijima Is., Okinawa (on *C. mydas* and *E. imbricata*); Yakushima Is., Kagoshima (on *C. caretta*); Tateyama, Chiba (on *C. caretta*); Otsuchi, Iwate (on *C. caretta*, and on *Chelonia mydas agassizii* [Hayashi *et al*. 2011]);

Hachijojima Is., Tokyo (on *C. caretta*); Hahajima Is., Ogasawara, Tokyo (on *C. mydas*); Minabe,

Wakayama (on *E. imbricata*); Motojima beach, Tanabe, Wakayama (on *C. mydas*); Manazuru, Kanagawa (on *C. mydas*); Fukui Prefecture (on *C. caretta* and *E. imbricata*); Niigata Prefecture (on *E. imbricata* [Hayashi 2021]). All other records are from Hayashi (2012).

**Habitat:** Found on carapace, plastron, head, flipper, limbs and soft skin of sea turtles.

***Platylepas ophiophila*** Lanchester, 1902 Fig. 2E-G

*Platylepas* -? Darwin, 1854: 430.

*Platylepas ophiophilus* Lanchester, 1902: 371, pl. 35 figs. 5-5b.

*Cryptolepas ophiophilus* Krüger, 1912: 12.

*Platylepas krügeri* Broch, 1931: 122.

*Platylepas ophiopholis* Nilsson-Cantell, 1938: 77.

*Platylepas indicus* Daniel, 1958: 755.

*Platylepas ophiophia* Ren, 1980: 191. – Liu & Ren, 2007: 316, fig. 140.

*Platylepas* sp. Hayashi, 2025: 65, fig. 2.

### Descriptions

Shell depressed, orifice large and ovoid. Overall shell outline variable, ranging from angular or sub-hexagonal to smoothly subcircular depending on the host substrate. Parietes aporous, externally marked with longitudinal ribs which are crossed by transverse grooves; however, the prominence of these external ornamentations varies significantly among individuals. The midribs on the rostrum and carina are a little shorter than those of the lateral compartments. Internally, the longitudinal ribs are visible in the lower half of the compartment, but the upper half of the shell is thickened considerably, growing inwards nearly to the level of the inner edge of the midrib. Basis membranous, adapting to the convexity of the host substrate. Opercular valves subequal, elongate and relatively narrow.

### Remarks

Importantly, our observations reveal that *P. ophiophila* exhibits extreme morphological plasticity depending on the host species. Specimens collected from sea snakes typically display the angular, sub-hexagonal shell outline and distinct surface ornamentation characteristic of historical descriptions (e.g., Lanchester, 1902; Utinomi, 1970). In stark contrast, specimens collected from the dugong lack these angular features and possess a smoother, subcircular shell. Morphologically, these sirenian-associated individuals are virtually indistinguishable from small specimens of *P. hexastylos*. This profound phenotypic plasticity highlights the limitations of relying solely on external shell morphology for species identification in *Platylepas*.

### Previous records on Japanese coast

Blue-banded sea snake, *Hydrophis cyanocinctus* Daudin, 1803, Mano Bay, Sado Island, Niigata, Japan (Utinomi 1970).

**Habitat:** attached to sea snakes and a dugong (new host record).

## MOLECULAR PHYLOGENY

The ML tree inferred from the concatenated COI and H3 sequences separated the examined *Platylepas* specimens into three clades corresponding to *P. decorata*, *P. hexastylos*, and *P. ophiophila* (Fig. 3A). With *Chelonibia caretta* used as the outgroup, *P. decorata* was placed outside the clade comprising *P. hexastylos* and *P. ophiophila*. The *P. hexastylos* clade included specimens collected from loggerhead, green, and hawksbill turtles, whereas the *P. ophiophila* clade included specimens collected from sea snakes and a dugong. Both the *P. hexastylos* and *P. ophiophila* clades were supported by high bootstrap values. Within *P. ophiophila*, the dugong- and sea-snake-derived specimens were arranged in two shallow subclades, each containing one specimen from each host type.

**Figure 3.**
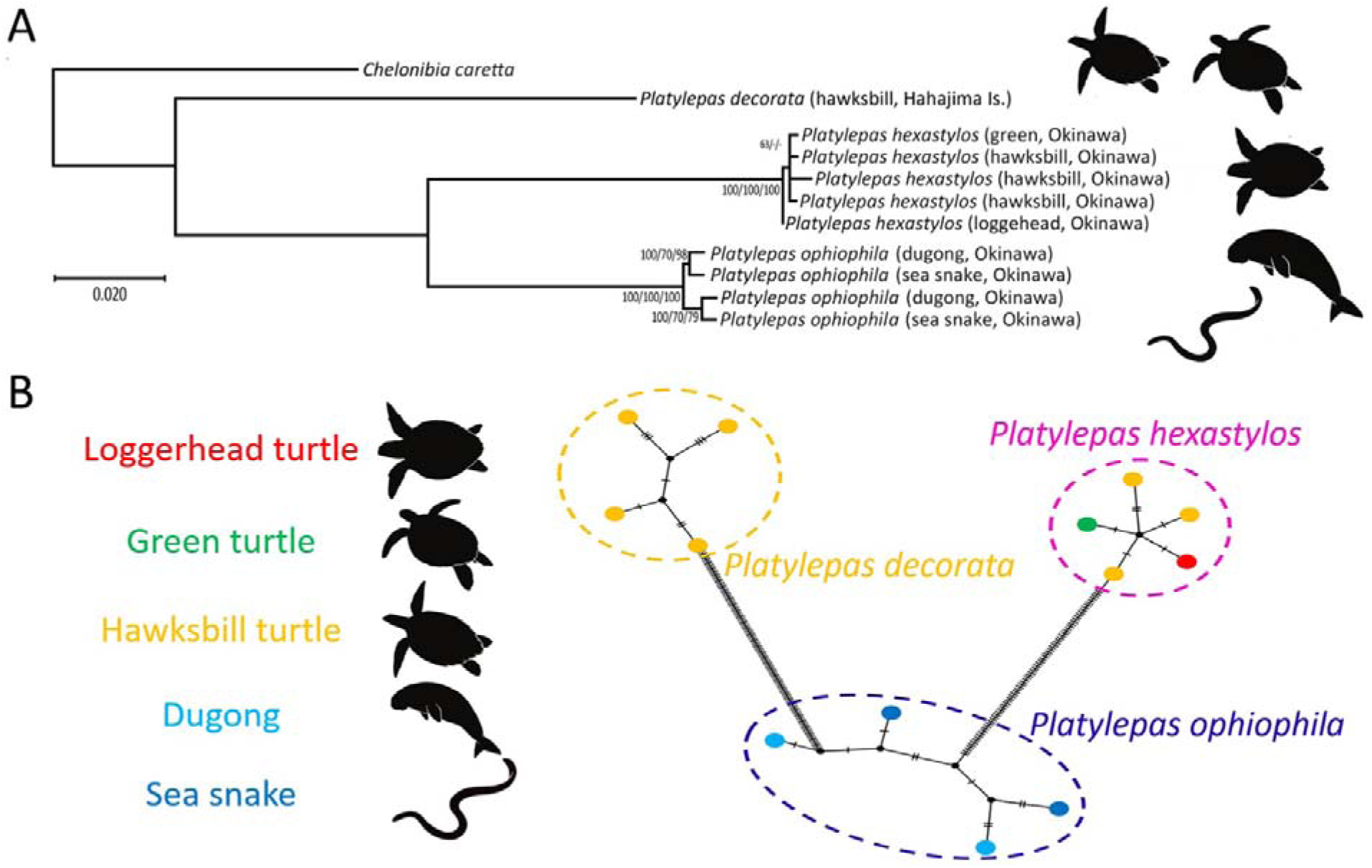
A. Molecular phylogenetic tree of the genus *Platylepas* and its outgroup, inferred using the Maximum Likelihood (ML) method based on concatenated COI and H3 gene sequences (781 bp). Numbers at the nodes, separated by slashes, represent bootstrap support values from ML, Neighbor-Joining (NJ), and Maximum Parsimony (MP) analyses, respectively (1000 replicates). The scale bar indicates the number of nucleotide substitutions per site. Silhouettes denote the host animal group for each major clade. **B.** COI haplotype network of *Platylepas* spp. TCS network (PopART) based on the 565-bp COI alignment; outgroups excluded. Each circle is a haplotype, with area proportional to sample size; colours indicate hosts as in the legend (loggerhead, green and hawksbill turtles; dugong; sea snake). Short hash marks on edges show single mutational steps; small black dots are inferred unsampled haplotypes (median vectors). Dashed loops outline the three species-level clusters (*P. decorata* n = 4; *P. hexastylos* n = 5; *P. ophiophila* n = 4). The two dugong-derived *P. ophiophila* samples fall into distinct haplotypes separated by few steps, consistent with intraspecific variation rather than cryptic species. Gaps were ignored (ε = 0); this figure is schematic and branch lengths are not to scale.

The COI haplotype network also separated the *Platylepas* specimens into three groups corresponding to *P. decorata*, *P. hexastylos*, and *P. ophiophila* (Fig. 3B). The *P. ophiophila* group contained haplotypes from both dugong- and sea-snake-derived specimens. The two dugong-derived *P. ophiophila* haplotypes were not identical and occupied different positions within the *P. ophiophila* network. Dugong- and sea-snake-derived haplotypes were connected within the same haplotype group by short mutational paths and did not form separate host-associated groups.

## DISCUSSION

### Morphological variation and species boundaries in *Platylepas*

A primary outcome of this study is that *P. ophiophila* is distinct from *P. hexastylos*, despite partial overlap in external shell morphology. Pfaller et al. (2012) suggested that barnacles found on sea snakes might represent a host-associated ecotype of the widespread turtle-associated species *P. hexastylos*. Our results do not support this interpretation. The concatenated COI and H3 phylogeny recovered *P. ophiophila* as a clade separate from *P. hexastylos*, and specimens from both sea snakes and a dugong were placed within this clade (Fig. 3A).

At the same time, our observations indicate that shell morphology in *P. ophiophila* is strongly influenced by host substrate. Sea-snake-associated specimens generally retain the angular shell outline and external ornamentation described in previous taxonomic accounts (Lanchester, 1902; Utinomi, 1970), whereas the dugong-associated specimens examined here are smoother and more subcircular, closely resembling small individuals of *P. hexastylos*. This host-associated morphological variation underscores the difficulty of identifying *Platylepas* species using external shell morphology alone. Species delimitation in *Platylepas* therefore requires an integrative approach that combines shell morphology, opercular characters, host association, and molecular data, a point that has also been discussed in relation to host-associated morphological variation in *Chelonibia* (Cheang et al. 2013; Zardus et al. 2014; Hyatt and Dunbar 2024).

### Ecological Specialisation and Implications for Host Movements

A host’s epibiotic fauna is intrinsically linked to its migratory behaviour and habitat selection. Unlike highly pelagic sea turtles, which often host broadly dispersed, panmictic barnacle populations (e.g., *P. hexastylos*), sea snakes tend to exhibit stronger site fidelity to specific coastal habitats, such as local seagrass beds (Lukoschek & Shine 2012; Udyawer et al. 2015; 2016).

Consequently, their primary epibiont, *P. ophiophila*, may develop more pronounced regional genetic structuring. In this study, the two *P. ophiophila* specimens derived from the same dugong fall on separate branches of the haplotype network (Fig. 3B); yet pairwise COI distances are well within intraspecific values and the H3 gene shows no concordant deep split (Table 2).

**Table 2.** Pairwise genetic distances (p-distance) in the genus *Platylepas* based on COI and H3 sequence variation (565 bp and 216 bp).

Notably, these two dugong-derived barnacles show genetic affinities to distinct *P. ophiophila* haplotypes recovered from sea snakes in geographically separate areas of Okinawa: Awase on the Pacific coast and Nakijin on the Motobu Peninsula (Fig. 1). This pattern is consistent with sequential recruitment from different nearshore habitats. One possible interpretation is that this host individual, potentially migrating from southern waters, accumulated distinct barnacle cohorts sequentially along the Okinawan coast—for instance, recruiting one haplotype while foraging near Awase, and acquiring another further north near Nakijin prior to its mortality.

The Okinawan dugong population is extremely small (Ozawa et al. 2024) and opportunities for direct tracking are limited. Because epibiotic barnacles can retain localized population structures that mirror the host’s passage through specific coastal water masses, phylogeographic analysis of these epibionts could serve as a powerful complementary tool to telemetry and stranding records for inferring host movement histories. Given the limited loci and small sample size analysed here, this remains a prospective application that requires validation with known-origin material and broader multilocus epibiotic sampling across multiple dugongs and sea snakes.

### Evolutionary history and host-use transitions in *Platylepas*

Having established the distinction between *P. ophiophila* and *P. hexastylos*, we next consider the evolutionary implications of host use in *Platylepas*. While *P. hexastylos* and *P. decorata* are epibionts of sea turtles—hosts that are highly migratory but also utilise seagrass beds—*P. ophiophila* appears to be a specialist on hosts more strictly dependent on seagrass ecosystems, namely sea snakes and dugongs. These two host groups, a reptile and a mammal, are taxonomically distant but are ecologically linked by their shared reliance on this specific habitat. Our phylogenetic analysis, in agreement with previous studies (Hayashi *et al*. 2013, Hayashi 2013b), demonstrates that *Platylepas decorata* and *P. hexastylos* do not form a monophyletic clade (Fig. 3A). Although the present study did not perform a molecular clock analysis, the branching order inferred from the COI and H3 markers recovered *P. decorata* as sister to the clade comprising *P. hexastylos* and *P. ophiophila*. This topology may suggest that the last common ancestor of these three sampled species was associated with sea turtles. This inference is consistent with fossil evidence, which is primarily based on ichnofossils (attachment scars) found on ancient turtle shells (Hayashi et al. 2013; Collareta et al. 2022).

A significant gap has long existed between the molecularly estimated origin of *Platylepas* in the Miocene and its body fossil record, which was, until recently, limited to the Pleistocene (Ross 1963, Mimoto 1991, Collareta *et al*. 2019, Karasawa & Kobayashi, 2022). Hayashi *et al*. (2013) hypothesised that this discrepancy could be resolved by identifying attachment scars on fossilised sea turtle carapaces. This proposition subsequently stimulated palaeontological research (Collareta *et al*. 2022; 2023; Perreault *et al*. 2025), which has indeed uncovered numerous such trace fossils, thereby strengthening the case for a deep evolutionary history of the genus in association with sea turtles. Our phylogeny provides further support for this narrative, showing a clear evolutionary trajectory that originated with turtle-associated ancestors.

The subsequent divergence of the *P. ophiophila* clade may represent a notable evolutionary transition associated with a change in host use from sea turtles to other inhabitants of seagrass ecosystems. A working hypothesis regarding the timing of this transition can be explored by considering published temporal frameworks for coronuloid evolution together with the fossil records of potential hosts (Fig. 4). Molecular-based estimates have suggested a Miocene timeframe for diversification within coronuloids/platylepadids (Hayashi et al. 2013), whereas some ichnofossil evidence (attachment scars) on fossil turtle shells has been interpreted to indicate that platylepadid-like associations could date back to the Oligocene (e.g., Collareta et al. 2022).

**Figure 4.**
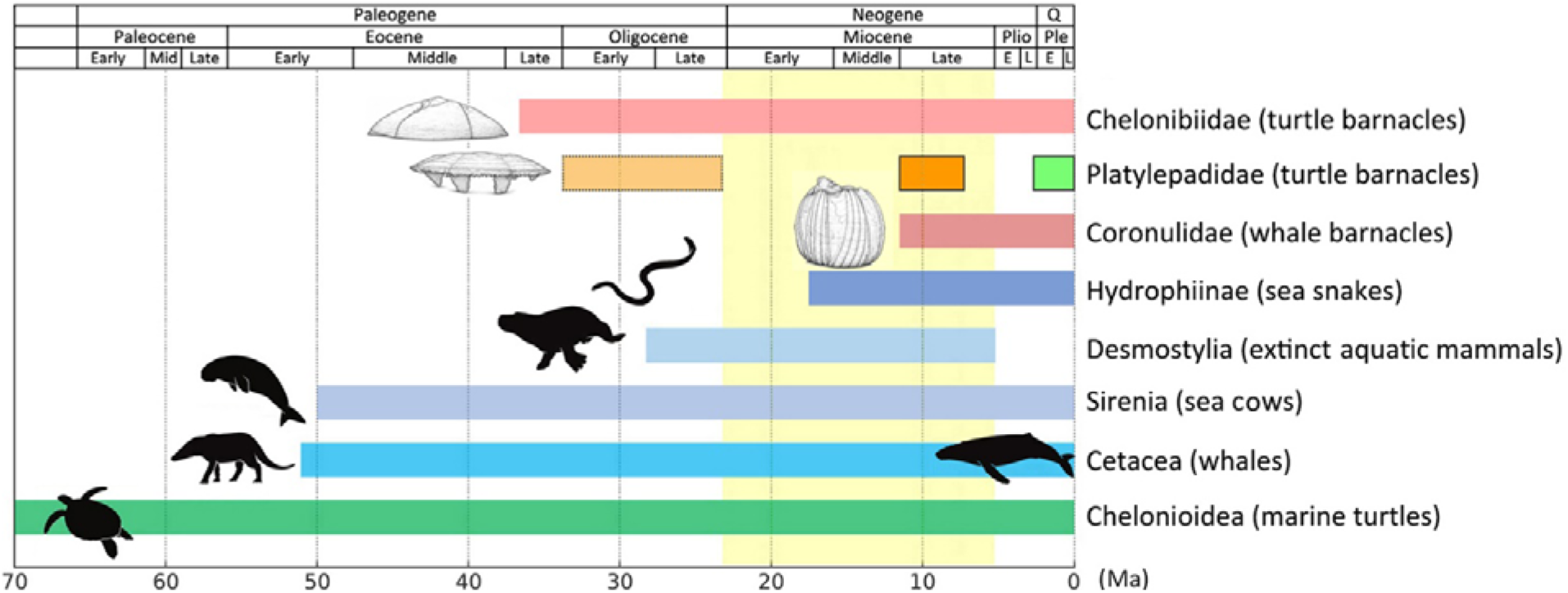
Schematic timeline showing temporal overlap among coronuloid barnacles and host clades. The pink bar indicates the oldest turtle barnacle linage (Chelonibiidae, including Chelonibiinae and Emersoniinae). The yellow band marks a Miocene window for the origin of *Platylepas* after Hayashi et al. (2013), the pale-yellow dashed block indicates Oligocene presumptive occurrences of platylepadids (Collareta *et al*. 2023), orange block indicates Miocene turtle ichnofossils attributable to platylepadids (Collareta *et al*. 2022), and the light green block indicates Pleistocene body-fossil record of platylepadids (Ross 1963, Mimoto 1991, Collareta *et al*. 2019, Karasawa and Kobayashi 2022). Red bar shows the predominantly Late Miocene body-fossil record of whale barnacles (Buckeridge *et al*., 2019). Horizontal bars denote broad geological ranges for sea snakes (Lee *et al*. 2016; Sanders *et al*. 2013), extinct aquatic mammals, Desmostylia (Asai *et al*. 2025), sea cows (Heritage and Seiffert, 2022), whales (Cabrera *et al*. 2021), and marine turtles (Cadena and Parham 2015).

If platylepadids were already present by the Miocene, this interval broadly overlaps with the diversification of early sirenians and the now-extinct desmostylians, large coastal herbivores thought to have inhabited seagrass or kelp-associated habitats. It is thus plausible that early host-use shifts within this lineage involved seagrass-associated marine mammals. In contrast, the main radiation of true sea snakes occurred later, from the late Miocene to Pliocene (Sanders et al. 2013). Taken together, these lines of evidence are consistent with (but do not demonstrate) a multi-stage colonisation scenario for the *P. ophiophila* lineage: an initial association with seagrass-linked mammals followed by a later expansion onto sea snakes as hosts became available in the same ecosystem (Fig. 4).

### Limitations and future work

Several issues remain to be resolved before species boundaries and host associations within *Platylepas* can be fully stabilised. First, the number of specimens analysed here is limited, particularly for *P. ophiophila* from sirenian hosts, and the molecular dataset is restricted to partial COI and H3 sequences. Broader sampling from sea snakes, sirenians, and sea turtles across the Indo-Pacific, combined with additional nuclear markers or genomic data, will be necessary to evaluate population structure, host specificity, and possible regional differentiation in greater detail.

Second, our discovery of *P. ophiophila* on a dugong indicates that historical records of *Platylepas* from sirenians require re-examination. Barnacles have previously been reported from dugongs in Moreton Bay, Australia (Darwin 1854), Magnetic Island, North Queensland (Zann and Harker 1978), Karratha, Nichol Bay, Northwestern Australia (Jones 2004), and New Caledonia (Fischer 1884), and these records were generally assigned to *P. hexastylos*. Because the dugong-associated specimens examined here are morphologically similar to small individuals of *P. hexastylos*, at least some historical sirenian-associated records may require reassessment using both morphology and molecular data.

Finally, the taxonomic status of *Platylepas coriacea* Monroe & Limpus, 1979, a putative leatherback turtle-associated species, remains an important unresolved issue in the taxonomy of *Platylepas*. This species is morphologically similar to *P. hexastylos*, and its validity should be reassessed through targeted comparisons of leatherback-associated specimens with *P. hexastylos* from other sea turtle hosts. Ideally, such reassessment should incorporate type or topotypic material where available, detailed morphological comparisons, and molecular data. A broader revision of *Platylepas*, including sirenian-, sea snake-, hard-shelled turtle-, and leatherback-associated material, will therefore be necessary to establish stable species boundaries and host-association patterns within the genus.

## ACKNOWLEDGEMENTS

We thank the fishermen of Toya fishery port for sea turtle epibiotic investigation, and the Nakijin Fisheries Cooperative Association and others who collected dead dugongs off the coast of Nakijin Village in Okinawa. We are grateful to Suma Aqualife Park (Kobe City) and the Okinawa Churaumi Aquarium (Okinawa Prefecture) for permissions and assistance related to sea snake husbandry and sampling, and the Seto Marine Biological Laboratory of Kyoto University for permission to reproduce the images of *Platylepas ophiophila* by Utinomi (1970) in Figure 2. The illustrations of marine vertebrates in Figures 1, 3 and 4 were made by Dr. Chihiro Kinoshita. The author used ChatGPT (OpenAI) and Gemini (Google LLC) to refine the English language and grammar of this manuscript. All authors reviewed and edited the output as needed and takes full responsibility for the content of the publication.

## CONFLICT OF INTEREST

The authors declare no conflict of interest.

## DATA AVAILABILITY

The materials examined were deposited in National Museum of Natural Science, Tokyo (NMNS-Cr-33017-33026) and all sequences were deposited in GenBank (Table 1).

## AUTHOR CONTRIBUTION

RH designed the research; oversaw specimen preparation and data collection; curated the data; performed analyses; prepared visualisations and was chiefly responsible for species identification. RH also wrote the first draft. RH, HO, TSe, TSa, and TK conducted field research and provided resources and samples; all authors contributed to coordinate refinement, review, and edit the final manuscript.

## FUNDING

This study was supported in part by the KAKENHI grant (JP19K04683 to RH and JP18J15154 to TSe) from the Japan Society for Promotion of Science.

## ETHICAL STANDARDS

The collection and keeping of sea snakes were conducted with the approval of the Mayor of Kobe City (Permit Kobe Sanitation No. 0715ZA0007, valid from March 2, 2016 to March 1, 2021) and the Governor of Okinawa Prefecture (Permit Okinawa-Tokuten No. 220, valid from February 15, 2023 to February 14, 2028), in accordance with the Permission for the Care and Keeping of Specified Animals under the Act on Welfare and Management of Animals.

## Notes

### Competing Interest Statement

The authors have declared no competing interest.

### Summary of Updates

This revised version updates the title, abstract, introduction, systematic accounts, molecular results, discussion, figures, and captions. The manuscript has been revised to clarify species boundaries and host associations in Platylepas, especially the distinction between P. ophiophila and P. hexastylos. The spelling of P. ophiophila has been revised and explained under the International Code of Zoological Nomenclature. The systematic section has been expanded with updated synonymies, descriptions, remarks, voucher information, and host records. The molecular phylogeny and COI haplotype network sections have been rewritten to clarify the placement of sea snake and dugong derived specimens. The discussion has been reorganized to address morphological variation, host movement inference from epibiotic barnacle haplotypes, host use evolution, and future work. Figures and captions have been updated accordingly.

